# Evolutionary relations between mycorrhizal symbiosis and plant–plant communication in trees

**DOI:** 10.1101/2022.04.04.486918

**Authors:** Akira Yamawo, Hagiwara Tomika, Satomi Yoshida, Ohno Misuzu, Riku Nakajima, Yusuke Mori, Tamayo Hayashi, Hiroki Yamagishi, Kaori Shiojiri

## Abstract

Ecological factors that drive the evolution of plant–plant communication via volatile organic compounds (VOCs) have not been elucidated. Here, we examined the relationship between type of mycorrhizal symbiosis (arbuscular mycorrhiza, AM; ectomycorrhizal mycorrhiza, ECM) and plant-plant communication within tree species. We hypothesized that ECM promotes plant-plant communication among conspecific individuals in trees, because it promotes their cooccurrence through positive plant-soil feedback. We tested communication using saplings of nine tree species with either AM or ECM, either exposed for 10 days to volatiles from an injured conspecific or not exposed. We evaluated the number of insect-damaged leaves and the area of leaf damage after 1 and 2 months in the field. Most exposed ECM-associated trees had less leaf damage than controls. However, AM-associated trees did not differ in leaf damage between treatments. We combined our results with those of previous studies and analysed the evolutionary relation between mycorrhizal type and the presence or absence of plant–plant communication within tree species. ECM symbiosis is associated with the evolution of plant–plant communication within species. These results suggest that the evolution of types of mycorrhizal symbiosis associates with the evolution of plant-plant communications within tree species.

## Introduction

Plant–plant communication mediated by volatile organic compounds (VOCs) has been studied in many species [1-3]. Damaged plants release specific VOCs such as green leaf volatiles and terpenoids. VOCs from herbivore-damaged plants activate the expression of resistance genes, prime resistance in surrounding undamaged plants [1,3-5], and alter the behavior of herbivores, deterring herbivory of undamaged plants [6-8]. Thus, plant–plant communication favours adaptability for received plants. However, some species do not show plant–plant communication. Although numerous studies focus on the mechanisms of plant– plant communication, it is unclear which ecological factors drive the evolution of plant–plant communication mediated by VOCs.

In general, plant–plant communication is considered to benefit species that grow at a high density, because the effects of VOCs reduce with distance. Plant–plant communication by volatiles was effective in a few species within about 10 m from a defoliated tree [9,10]. These findings suggest that plant–plant communication evolved in association with population density.

Plant–soil feedback (PSF), mediated through interaction with microbial communities, regulates tree population density. The direction (positive or negative for conspecific individuals) of PSF effects depends on the type of mycorrhizal symbiosis. Bennett et al. [11] showed that arbuscular mycorrhizal (AM)-associated trees had negative effects on growth or survival of conspecific seedlings (negative soil–plant feedback), whereas ectomycorrhizal (ECM)-associated trees had positive effects (positive soil–plant feedback). These PSF effects correlated with population density in the field. Thus, mycorrhizal type and plant–plant communication may have evolved together.

We hypothesized that ECM symbiosis, which is positively associated with tree population density, promotes communication between conspecific individuals (Fig. 1). We conducted an experiment in two study fields using nine tree species with either AM or ECM symbiosis (Table S1). Thirty saplings of each species were exposed for 10 days to conspecific neighbours that had been damaged by clipping. Another 30 saplings were not exposed as controls. All saplings were then set in natural fields, and we evaluated the number of insect-damaged leaves at 1 month and the area of leaf damage at 2 months. We also compared two plant defence hormones, jasmonic acid (JA) and salicylic acid (SA), between treatments at 10 days.

**Fig. 1.**
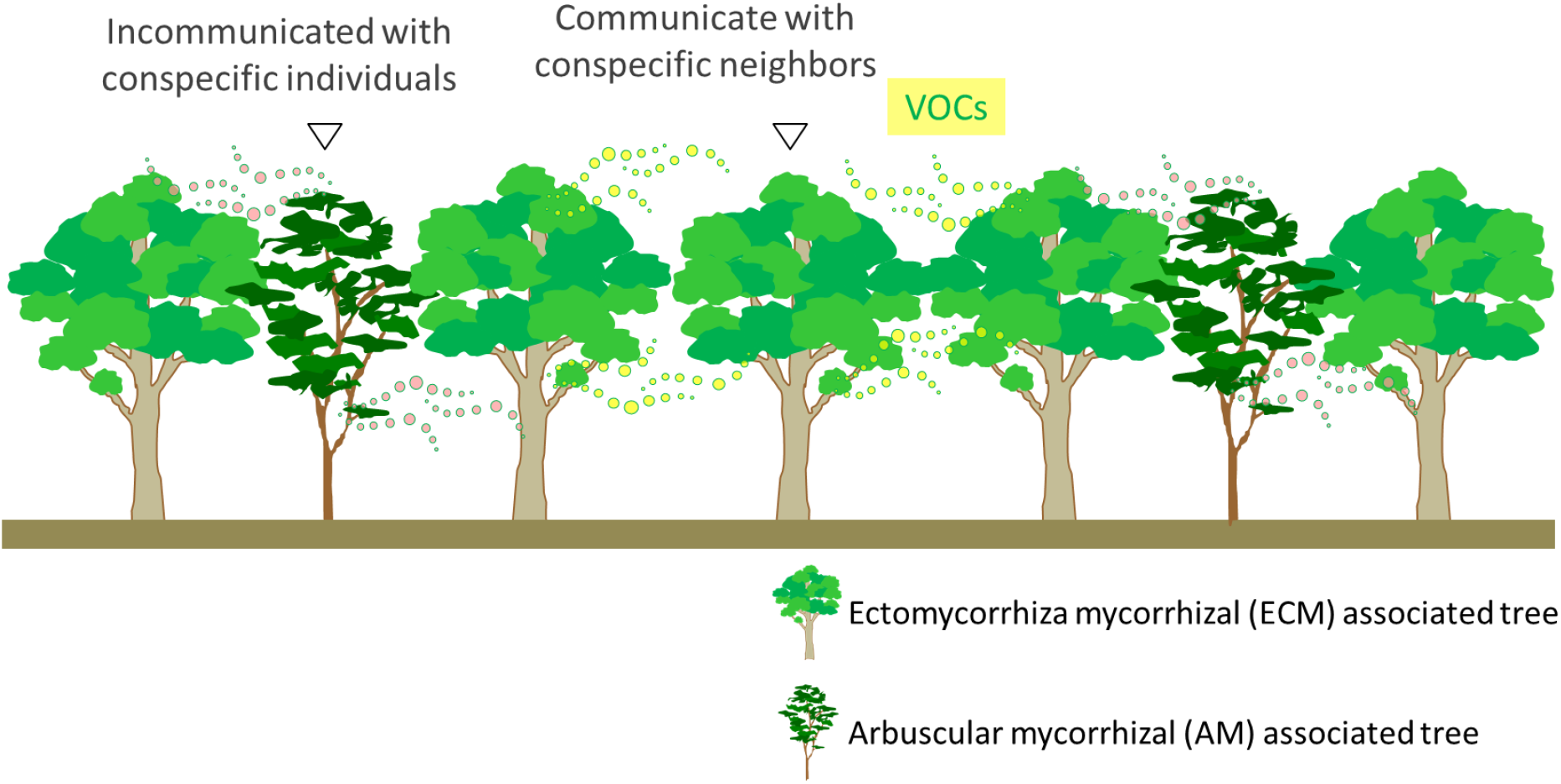
Conceptual diagram of our hypothesis. ECM symbiosis, which is positively associated with tree population density, promotes communication between conspecific individuals.

## Materials and methods

### Experimental species

We used nine plant species: five AM-associated species (*Magnolia obovata, Cerasus jamasakura, Viburnum dilatatum, Viburnum furcatum*, and *Aesculus turbinata*) and four ECM-associated species (*Pinus densiflora, Betula platyphylla, Quercus crispula*, and *Quercus serrata*) [12-16]. The natural population densities of ECM-associated species are higher than those of AM-associated species (Table S1) [17].

Because plant defenses for future growth are often developed at the seedling stage [18], we purchased sixty 3- to 5-year-old saplings of each species from a local nursery in Fukuoka City in November 2020. Each sapling had no large lateral branches, and a height of 40 to 50 cm. *Betula platyphylla* is known to communicate with conspecific neighbours through VOCs [19].

### Plant culture

In April and May 2021, all saplings were individually planted in plastic pots (15 cm × 15 cm × 25 cm) containing 50% tuff loam and 50% humus. The pots were maintained in a greenhouse at Hirosaki University (40°59’
sN, 140°47’E) or Ryukoku University (34°96’N, 135°93’E). The experiment began when new leaves reached the three- or four-leaf stage, in early spring (April), when plant defences for future growth often develop [20].

### Experimental design

Saplings of each species were randomly assigned to either the control (no exposure to VOCs, *n* = 30) or the exposure treatment group (exposed to VOCs, *n* = 30). The two groups were held in different area more than 10m away and separated by a glass wall at each university. These places were concrete ground and there were no plants around. In the exposure area, we placed five conspecific “emitter” plants on which half of every leaf was clipped, sited 60 cm apart. Around each emitter plant we arranged six experimental plants 30 cm away. In the control area, the experimental plants were placed around five non-damage plants. After 10 days, all plants were set in natural conditions of each tree species (The Shirakami Natural Science Park, Hirosaki University (40°52’
sN, 140°22’E) (*M. obovate, V. dilatatum, A. turbinate, B. platyphylla, Q. crispula*); or Ryukoku University Forest (34°96’N, 135°94’E) (*Cerasus jamasakura, Viburnum wrightii, Pinus densiflora, Quercus serrata*)) for 2 months. This site was an open land, surrounded by deciduous broad-leaved trees and pine trees, including experimental tree species. Pots were placed randomly 60 cm apart. The plants were watered every 2 days.

### Evaluation of defence induction

As indicators of defence induction, we counted the number of insect-damaged leaves at 1 month and measured the area of leaf damage at 2 months after treatments. We also compared two plant defence hormones, jasmonic acid (JA) and salicylic acid (SA), between treatments at 10 days after treatment.

### Number of damaged leaves and area of leaf damage

In all species except *P. densiflora*, at 1 month, on 10 or 12 September 2021, we measured herbivory by counting the leaves with any visible damage caused by insect herbivores on assay branches in both treatments. We have used this presence/absence measure of herbivory in our many previous works and found that it correlates with the percentage of leaf area removed. In *Pinus densiflora*, we counted the leaves damaged had turned brown marks as leaf damage by stink bugs, because they had no leaf-area loss and had only dark stab marks.

At 2 months, all leaves were collected and scanned on an image scanner (PM-850; Seiko Epson Corp., Suwa, Japan). The area of each leaf was subsequently measured in Scion Image photo-image analysis software (Scion Image; Scion Corp., Frederick, MD, USA). The area that had been consumed by herbivores was estimated by comparison with an uninjured leaf of equal size. *Cerasus jamasakura* was excluded from this analysis because of a technical mistake when the leaves were collected. In *Pinus densiflora*, we counted the leaves damaged by stink bugs per randomly selected 500 leaves, because they had no leaf-area loss and had only dark stab marks.

### Plant hormones

To determine any response induced in leaves exposed to VOCs, we analysed the contents of JA and SA in leaves by liquid chromatography – tandem mass spectrometry (LC-MS) according to Ozawa et al. [21] in 1–2 mm of leaf cut from each plant with scissors after 10 days’ exposure in both treatments.

Leaves (ca. 0.5 g) were immediately frozen in liquid nitrogen, homogenized in ethyl acetate (2.5 mL), and spiked with 10 ng of d2-JA (Tokyo Chemical Industries Co., Tokyo, Japan) and 1 ng of d4-SA (C/D/N Isotopes, Pointe-Claire, QC, Canada) as internal standards. After centrifugation of the mixture at 2300× *g* for 10 min at 4 °C, 1 mL of supernatant was transferred to a 1.5-mL tube and then evaporated to dryness under vacuum. The residue was suspended in 50 µL of 70% methanol/water (v/v) and centrifuged, and the supernatant was analysed by LC-MS/MS (LCMS-8050, Shimadzu, Kyoto, Japan). Analytes were separated by high-performance liquid chromatography through a Mightysil RP-18 GP column (100 mm × 2.0 mm, 3 μm particle size, Kanto Chemical, Tokyo, Japan) at a flow rate of 200 µL min^−1^ with a linear gradient (0.1% formic acid aq. “A” and methanol “B”; 5%–95% B/(A + B) for 16 min). Concentrations of JA, d2-JA, SA, and d4-SA were determined by multiple reaction monitoring (MRM). The monitored mass transitions were *m/z* 209 to *m/z* 59 for JA, *m/z* 211 to *m/z* 59 for d2-JA, *m/z* 137 to *m/z* 93 for SA, and *m/z* 141 to *m/z* 97 for d4-SA. The conditions for MS were optimized for MRM with authentic d2-JA and JA (Tokyo Chemical Industries), d4-SA (C/D/N Isotopes), and SA (Wako Pure Chemical Industries, Osaka, Japan). Samples with technical mistakes were removed from all data analysis.

### Data analysis

All data were analysed in R v. 4.1.0 software [22]. All data met the statistical assumptions of normality and homoscedasticity according to the Kolmogorov–Smirnov test and *F*-test, and analyses performed depended on the dataset structure. All tests were two tailed, with *P* < 0.05 considered significant.

### Number of damaged leaves, area of leaf damage, and plant hormones

Number of damaged leaves and area of leaf damage were analysed by general linear models (GLMs) with Poisson or Gamma distributions, and tested by chi-squared test. Damage of *P. densiflora* by stink bugs was analysed by a GLM with a Poisson distribution, and tested by chi-squared test. Number of damaged leaves, area of leaf damage or damage by stink bugs were included as response variables, and exposure treatment, mycorrhizal types and their interaction were included as explanatory variables. Total number of leaves was included as a covariate.

Concentration of plant hormones were compared between control and voc-exposed plants by a GLM with a negative-binomial distribution, and tested by chi-squared test. A false discovery rate correction for multiple comparisons was then applied.

### Phylogenetic correlation between mycorrhizal types and plant–plant communication

We searched studies in Google Scholar with the ’Plant-plant communication’ and ’tree’. We also data on presence or absence of plant-plant communication were collected from the list by Heil & Karban [1]. Our database contains information on the mycorrhizal types, and plant-plant communication of 21 species in 12 genera in 8 families. To prepare a phylogenetic tree, species names were standardized against Plant List 1.1 (http://www.theplantlist.org/), and membership in higher taxonomic groups was standardized against APG IV [23].

The phylogenetic tree was constructed in PhytoPhylo tree software [24], being based on the phylogeny generated by Zanne et al. [25] and updated by Qian and Jin [24]. The final phylogenetic tree had 21 tip labels and 19 internal nodes in scenario 3, the accuracy of which has been verified (Fig. S1).

The PhytoPhylo phylogenetic tree was used for phylogenetic generalized least squares (pGLS). We used Pagel’s λ correlation structure [26] to incorporate the phylogenetic information into the models. The models were constructed by using the corPagel function in the “ape” package [27] with the amount of phylogenetic signal in the data estimated by the restricted maximum likelihood approach [28]. The presence or absence of plant–plant communication was included as a response variable, and mycorrhizal type was included as an explanatory variable.

## Results and Discussion

Effects of exposure treatment on leaf damage differed clearly among mycorrhizal types at both 1 month (χ^2^ = 81.10, *P* < 0.001; Fig. 2) and 2 months (χ^2^ = 14.89, *P* < 0.001; Fig. 3). Both number of damaged leaves and area of leaf damage in exposed ECM species—*Betula platyphylla, Quercus crispula* and *Q. serrata* (but not *Pinus densiflora*)—were significantly smaller than those in control trees (1 month, χ^2^ = 93.06, *P* < 0.001; 2 months, χ^2^ = 52.01, *P* < 0.001). However, they did not differ between treatments in AM species, *Magnolia obovata, Cerasus jamasakura, Viburnum dilatatum, Viburnum wrightii*, and *Aesculus turbinata* or *Pinus densiflora* (AM trees: 1 month, χ^2^ = 0.86, *P* = 0.35; 2 months, χ^2^ = 0.34, *P* = 0.56; *P. densiflora*: 1 month, χ^2^ = 0.23, *P* = 0.63; 2 months, χ^2^ = 1.30, *P* = 0.25).

**Fig. 2.**
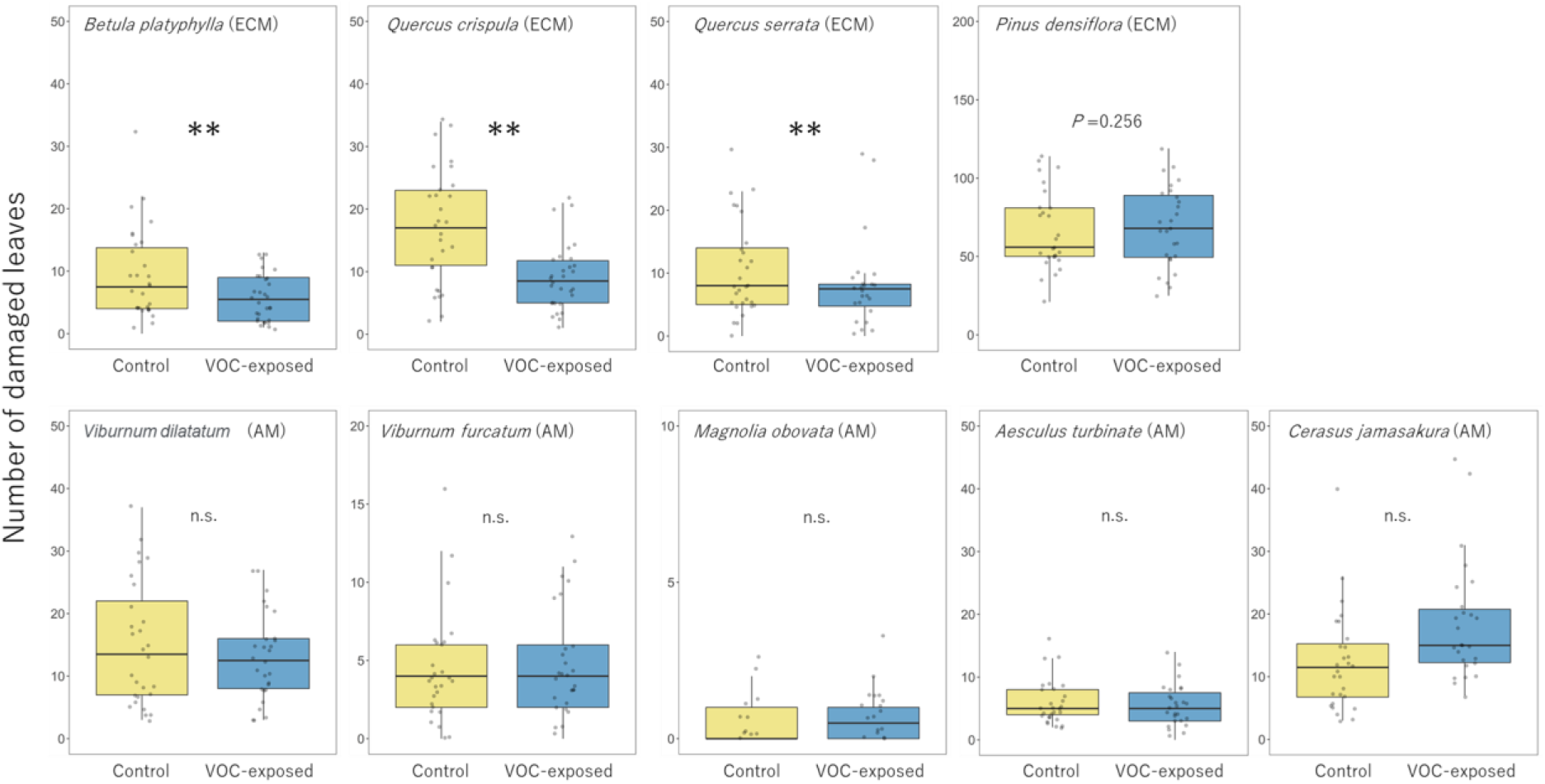
Number of damaged leaves at 1 month in control and exposed plants. Values in *Pinus densiflora* are the number of damage marks caused by stink bugs. ** indicate *P* < 0.05.

**Fig. 3.**
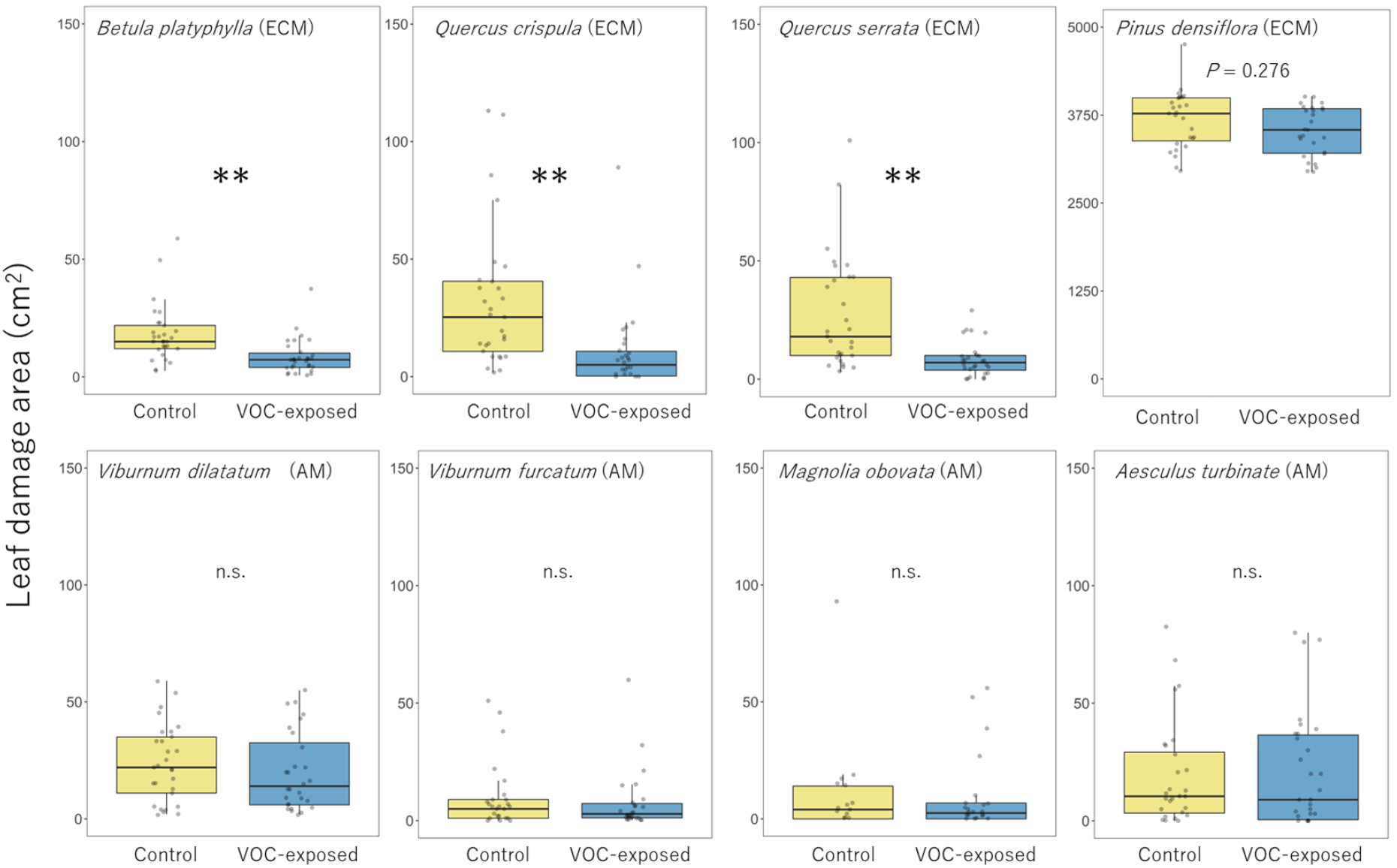
Area of leaf damage at 2 months in control and exposed plants. Values in *Pinus densiflora* are the number of damage marks caused by stink bugs. ** indicate *P* < 0.05.

The plant hormone assays also showed evidence of plant–plant communication in two species. Exposure increased JA in *B. platyphylla* (Table S2), supporting the results of reduced leaf damage. On the other hand, we could not find differences in plant hormones in *Q. crispula* or *Q. serrata*, despite these species’ significant differences in leaf damage between treatments. This discrepancy suggests that the defence response was induced after the hormone measurements (10 days from start of exposure treatment) in *Quercus*. In addition, JA and SA contents in *C. jamasakura* were significantly higher in exposed trees, despite no difference in leaf damage (Table S2). This suggests that induced responses in *C. jamasakura* are too small to affect leaf damage by insect pests. JA and SA contents did not differ between treatments in other species. These results highlight the importance of carefully assessing plant–plant communication in a variety of ways and differences in induction timing.

Overall, our experiment revealed that plant–plant communication between conspecific individuals was more common in ECM-associated tree species. Some plants release specific VOCs against specific herbivores [2,29]. However, as we used artificial clipping by scissors, our experiment reveals interspecific variations in plant–plant communication in response to common leaf damage such as by generalist herbivores.

Finally, we analysed the evolutionary relations between mycorrhizal type and presence or absence of plant–plant communication within species. Combining our results with previously published data, we found marginal or significant phylogenetic correlations of type of mycorrhizal symbiosis with plant–plant communication (*C. jamasakura*e include as non-communicated: λ = 1, *F* = 15.76, *P* < 0.001; *C. jamasakura* include as communicated: λ = 1, *F* = 3.53, *P* = 0.075). Although our experiment and analysis include only 21 tree species, these results suggest that mycorrhizal symbiosis has correlated the evolution of plant–plant communication within species.

Several studies reported that within-species PSFs differ between AM- and ECM-type trees [30]. For ECM-associated trees, PSFs are positive within species: growth and survival are increased in soil in which conspecifics grow [11]. Conversely, for AM-type trees, PSFs are negative within species: growth and survival are reduced in soil in which conspecifics grow [11]. The effects of these different PSFs promote variation in population density in tree species [11]. Yamawo & Ohno [31] found that ECM types are more likely to evolve short-distance seed dispersal. This is a positive PSF effect of ECM symbiosis. The evolution of short-distance seed dispersal should promote high population density and may promote the evolution of plant–plant communication. The presence/absence of plant–plant communication within our nine tree species was positively correlated with population density (*C. jamasakura*e include as non-communicated: χ^2^ = 11.5, *P* < 0.001; *C. jamasakura*e include as communicated: χ^2^ = 7.3, *P* < 0.001). Thus, differences in PSFs and population density may drive the evolution of plant–plant communication. This hypothesis should be tested in future studies of the evolution of plant–plant communication in trees.

Another possible explanation of correlation between mycorrhizal types and plant-plant communication is physiological differences among plants in association with mycorrhizal type. AM and ECM provide different ecological and physiological states for their host trees [32]. For example, AM extracts soil P at higher efficiency than ECM [33]; and ECM may have a greater symbiotic cost than AM [34,35]. These differences may underlie differences in the physiological conditions of trees such as amounts and compositions of VOCs. However, many AM-associated herbs and shrubs also communicate [1,2]. This fact suggests that AM physiology does not inhibit plant–plant communication, but instead population density explains the link between mycorrhizal type and the evolution of communication in trees. In any mechanisms, it seems likely that mycorrhizal symbiosis has driven the evolution of plant– plant communication in trees.

Plants have lived in close association with AM fungi for over 400 million years [30]. Most ECM-type trees evolved in some clade in the Cretaceous (between 145 and 66 Mya) [30]. If ECM symbiosis promoted the evolution of plant–plant communication, this may have happened in the Cretaceous. We do not know whether these ancient ECM-type trees had positive PSFs on conspecific individuals, but if they had, PSFs may have driven the evolution of plant–plant communication. This raises a novel question: at what point in evolutionary time did PSFs appear and how did they contribute to plant–plant communication? This question may be clarified by elucidating the mechanisms of PSFs in detail and mapping the evolutionary history of the microbes involved.

## Author contributions

Akira Yamawo (AY) and Kaori Shiojiri (KS) developed the core idea. AY, KS, Hagiwara Tomika (HT), Riku Nakajima (RN), Yusuke Mori (YM), Tamayo Hayashi, Satomi Yoshida and Ohno Misuzu (OM) performed field experiments. HT, RN and YM measured plant hormones. Hiroki Yamagishi supported all field experiments. AY and OM performed phylogenetic analysis. AY, HT and KS analysed data and wrote the paper. All authors contributed critically to the drafts and gave final approval for publication.

## Acknowledgements

We wish to thank Shirakami Research Center for Environmental Sciences of Hirosaki University for fieldwork assistance. We thank Haruna Ohsaki, Riku Nomiya, Taisei Kodera, Riki Iwabuchi, Nanaha Mizutani, Yuuta Dote, Ryota Kamidate and Hiyori Sasaki for their support during the experiment. This work was supported by a Japan Society for the Promotion of Science (JSPS) Grant-in-Aid for Scientific Research (KAKENHI; no. 19H03295 to AY) and by the Sumitomo Foundation’s Fiscal 2019 Grant Basic Science 301 research projects (no. 190876 to KS).

## Data availability statement

All datasets are available from the corresponding author on reasonable request.

## Code availability statement

The computer code is available from the corresponding author on reasonable request.

## Figure legends

**Fig. S1.**
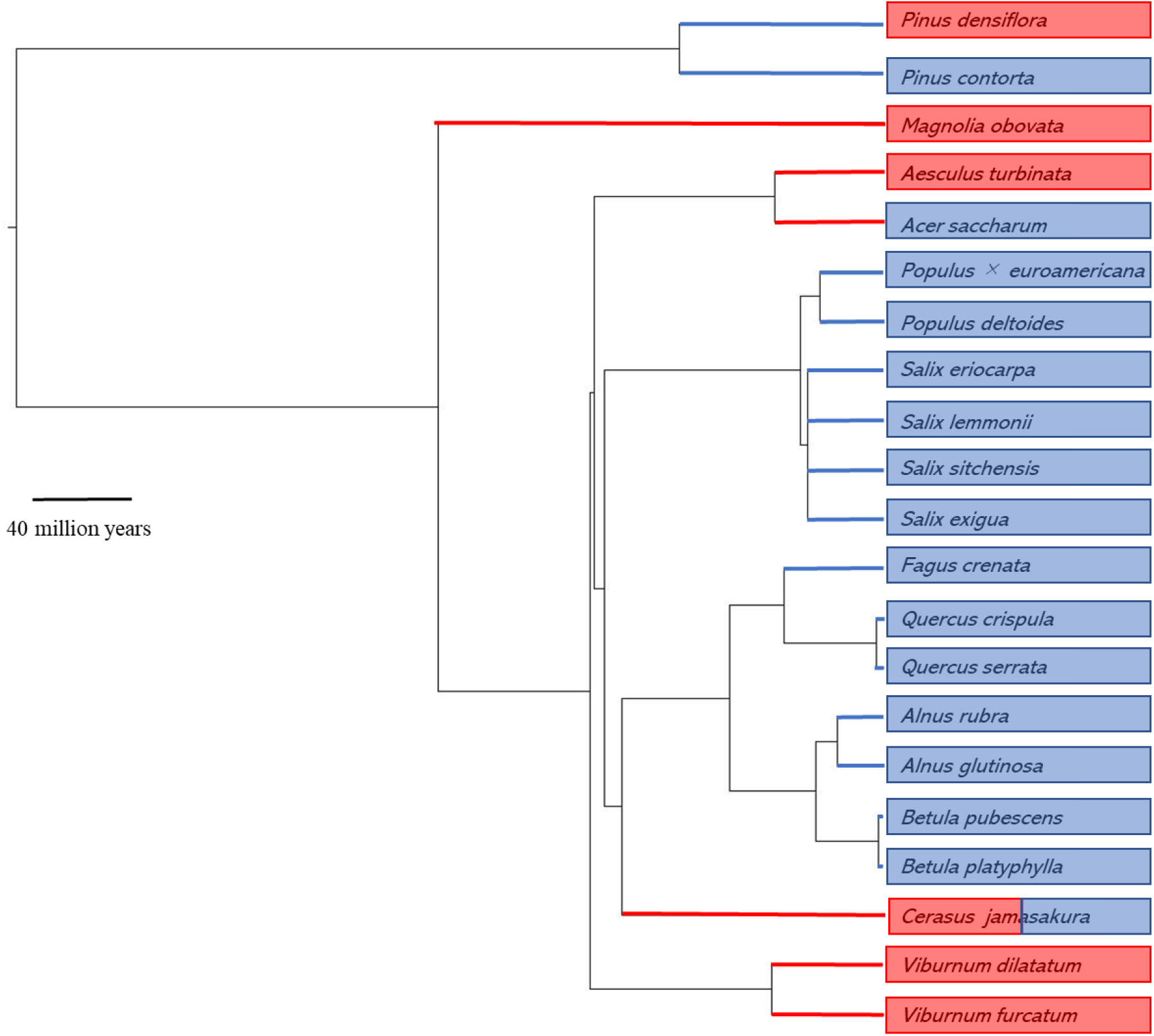
Time-calibrated phylogeny of the 21 plant species used in this study and previous studies (supplementary data) provided by Scenario 3 of S.PhyloMaker. Branch colour denotes mycorrhizal type (red, AM; blue, ECM). Colours of inner and outer labels ringing the tree indicate communication ability (red, absence; blue, presence). In *Cerasus jamasakura* is coloured in both red and blue, because this species had different results between herbivory and plant hormones-based estimation. The scale bar represents 40 million years.

**Table S1.** Population densities used in study fields.

| Study species | Mychorrhizal type | n | Population density (/10,000m<sup>2</sup>) |
| --- | --- | --- | --- |
| *Aesculus turbinata* | AM | 5 | 30.2 ± 19.1 |
| *Magnolia obovate* | AM | 19 | 23.0 ± 32.9 |
| *Cerasus jamasakura* | AM | 10 | 30.4 ± 18.7 |
| *Viburnum furcatum* | AM | 6 | 28.2 ± 31.1 |
| *Viburnum dilatatum* | AM | 1 | 2 |
| *Betula platyphylla* | EcM | 6 | 84.9 ± 124.1 |
| *Pinus densiflora* | EcM | 2 | 36.0 ± 31.1 |
| *Quercus crispula* | EcM | 3 | 80.0 ± 131.63 |
| *Quercus serrata* | EcM | 15 | 162.9 ± 358.0 |

*n*: number of sites.
Data are based on the “Monitoring Sites 1000” program of nationwide long-term monitoring of tree communities
in Japan [17].

**Table S2.**
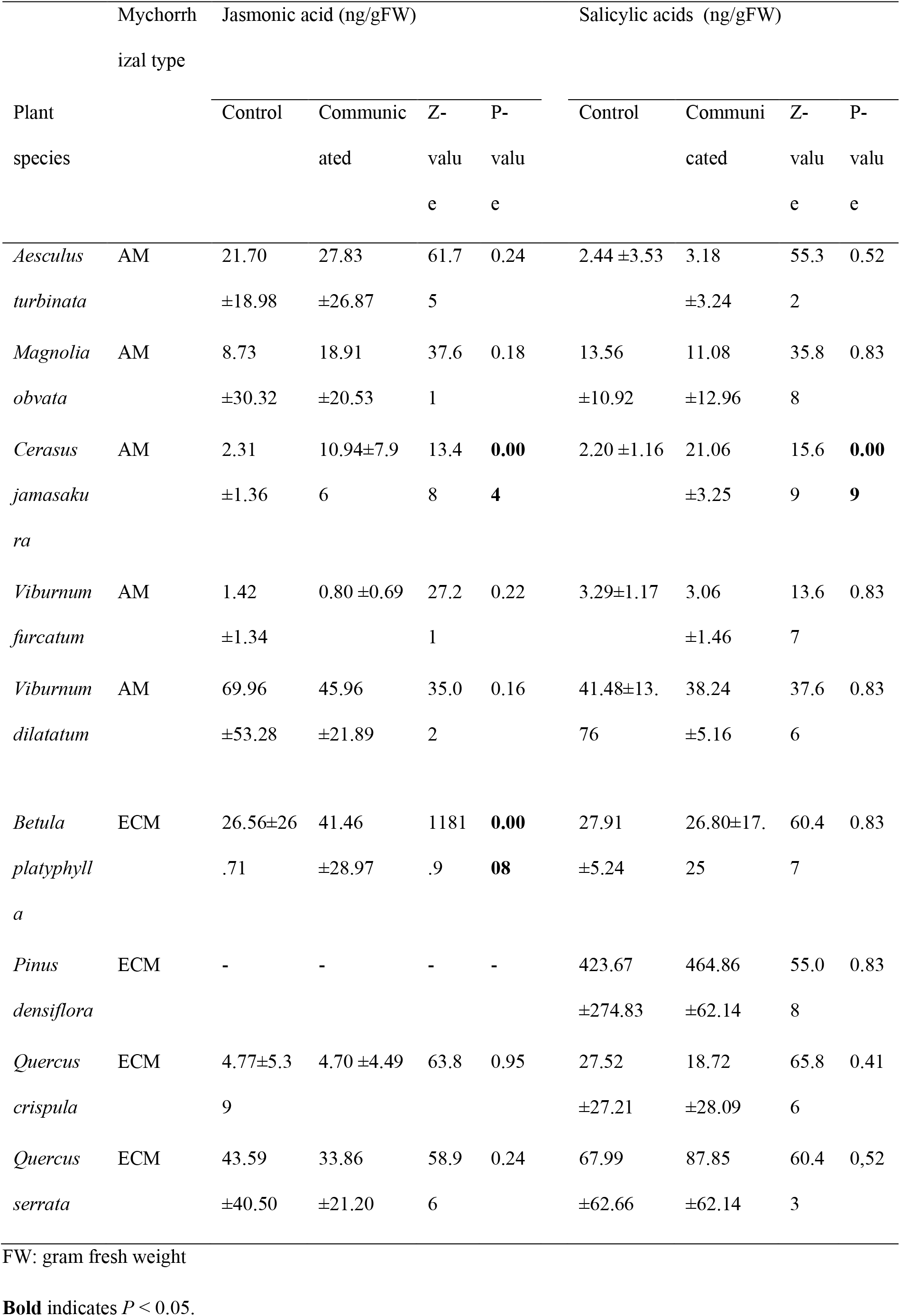
Contents of plant defence hormones in control and exposed plants.

